# Dual action of sphinganine in the plant disease resistance to bacteria

**DOI:** 10.1101/2024.02.07.579277

**Authors:** Eloïse Huby, Sandra Villaume, Catherine Chemotti, Stéphan Dorey, Sylvain Cordelier, Jérôme Crouzet, Guillaume Gilliard, Christine Terryn, Alexandre Berquand, Cornelia Herrfurth, Ivo Feussner, Cédric Jacquard, Florence Fontaine, Christophe Clément, Fabienne Baillieul, Magali Deleu, Sandrine Dhondt-Cordelier

## Abstract

Sphingolipids are ubiquitous, highly diverse molecules constituting at least 40% of plant plasma membranes. Initially known as modulators of membrane integrity, they now emerge as important players in plant responses to (a)biotic stresses. The interaction between *Arabidopsis thaliana* and the bacterium *Pseudomonas syringae* pv. *tomato* DC3000 *AvrRpm1* (*Pst AvrRpm1*) culminates in the activation of a programmed cell death known as the hypersensitive response, which is part of the plant immune response. In this study, we showed that the co-infiltration of *Pst AvrRpm1* and sphinganine (d18:0) in *Arabidopsis* leaves suppress the hypersensitive response. This suppression phenotype is also observed with bacteria carrying the effectors AvrB and AvrPphB but not with the ones carrying AvrRpt2 and AvrRps4. Sphingolipid-induced hypersensitive response suppression by *Pst AvrRpm1* is correlated with the down-regulation of the gene *AtNMT1* encoding a *N*-myristoyltransferase. d18:0 does not have a direct antibacterial effect and its co-infiltration in plants does not display typical signs of immune response such as activation of salicylic acid signaling pathway and extracellular reactive oxygen species production. Biophysical studies showed that d18:0 interacts with plant plasma membrane lipids. More specifically, d18:0 disturbs plant plasma membrane organization and mechanical properties. Our results demonstrate that sphingolipids play an important role in plant resistance, especially by interfering with the plasma membrane organization and effector localization and thus disturbing their function and subsequent immune responses.

## INTRODUCTION

Recognition of pathogenic microorganisms leading to plant immunity is commonly divided into two pathways that can mutually potentiate each other (Ngou *et al*., 2021). Pathogen perception involves the recognition of pathogen-associated molecular patterns (PAMPs) through plant cell surface-anchored pattern recognition receptors (PRRs), which induces pattern-triggered immunity (PTI) (Saijo and Loo, 2020). However, some bacteria can interfere with PTI by delivering effectors into the plant cell to block signaling mechanisms at different steps (Desveaux *et al*., 2006; Kamoun, 2006; Dodds *et al*., 2009). The recognition of microbial effectors or virulence factors by the plant cells through intracellular nucleotide-binding leucine-rich repeat receptors (NLRs) induces effector-triggered immunity (ETI). PTI and ETI share many signals and components, but their signaling pathways are induced at different scales and time (Bjornson and Zipfel, 2021; Kadota *et al*., 2019; Yuan *et al*., 2021). Typical overlaps in signaling components are e.g., the production of reactive oxygen species (ROS), the influx of calcium, or mitogen-activated protein kinase (MAPK) signaling cascades. They also share some immune responses, such as transcriptional reprogramming and alterations in stress-related hormone homeostasis, including salicylic acid (SA), jasmonic acid (JA), and/or ethylene (ET) (Bari and Jones, 2009; Betsuyaku *et al*., 2018; Guan *et al*., 2015). Rapid programmed cell death (PCD) at the site of infection, known as the hypersensitive response (HR), usually is a hallmark of the ETI pathway (Couto and Zipfel, 2016; Henry *et al*., 2013; Kadota *et al*., 2019; Ngou *et al*., 2021; Yuan *et al*., 2021).

The bacterial pathogen *Pseudomonas syringae* pv. *tomato* DC3000 (*Pst* DC3000) has been extensively studied due to its capacity to cause bacterial speck in both tomato and *Arabidopsis thaliana* (hereafter *Arabidopsis*) (Lee *et al*., 2015; Wei *et al*., 2018), as well as the availability of its complete genome sequence (Buell *et al*., 2003). Transformation of *Pst* DC3000 to express effectors has been essential for the discovery of mechanisms associated with ETI (Boyle and Martin, 2015; Hicks and Galán, 2013; Nimchuk *et al*., 2000). AvrRpm1 effector was originally identified in *P. syringae* pv. *maculicola* strain m2 (Debener *et al*., 1991) and harbors an N-terminal eukaryotic consensus site for myristoylation required for its full avirulence function. This allows the effector, once acylated by the plant host, to be redirected to the plant plasma membrane (PM) (Boyle and Martin, 2015; Hicks and Galán, 2013; Nimchuk *et al*., 2000). Although the host acylation apparatus is initially exploited by bacteria to interfere with PTI by binding to the kinase domain of PRRs (Boyle and Martin, 2015), in resistant plants, it can actually turn against the pathogen. For instance, the NLR RPM1 (RESISTANCE TO *PSEUDOMONAS SYRINGAE* PV *MACULICOLA*) in *Arabidopsis*, is monitored by the PM-associated protein RPM1-INTERACTING PROTEIN-4 (RIN4), which guards it against effector-mediated perturbations (Boyes *et al*., 1998; Jones and Dangl, 2006; Mackey *et al*., 2002). Therefore, once relocated to the PM, AvrRpm1 will target RIN4, inducing its phosphorylation. This activates RPM1 which triggers the ETI and the HR, together interrupting the bacterial spread of the and leading to the degradation of RPM1 (Boyes *et al*., 1998; Mackey *et al*., 2002; Xin *et al*., 2018).

Sphinganine is a sphingolipid (SL), belonging to the long-chain base (LCB) class, which are one major building block of SLs (Pata *et al*., 2010). It consists of a C18 core with an amino group at C2 and two hydroxylations at C2 and C3 (Luttgeharm *et al*., 2016; Mamode Cassim *et al*., 2019). SLs are key components of the plant PM structure, accounting for up to 40 mol% of its external leaflet (Cacas *et al*., 2016). They are found mostly in lipid microdomains, and play an important part in signaling, protein aggregation, and stress response (Cacas *et al*., 2016; Grennan, 2007; Mamode Cassim *et al*., 2019; Pata *et al*., 2010). Recent studies have shown that SLs are key players in PCD regulation occurring either during plant development or immunity (Ali *et al*., 2018; Berkey *et al*., 2012; Huby *et al*., 2020; Lenarčič *et al*., 2017). For instance, Bax-inhibitor-1 interacts with enzymes of the SL biosynthetic pathway FAH1 and 2 (FATTY ACID HYDROXYLASE 1 and 2). It leads to changes in lipid content of membrane microdomains (Lenarčič *et al*., 2017). This causes the loss of proteins normally located in these microdomains and which are essential for plant defense, especially for cell death by oxidative stress or SA (Ishikawa *et al*., 2015). Further links between PCD, sphingolipids and the SA pathway have recently been highlighted. In *Arabidopsis*, triple mutations of enzymes of the SL biosynthetic pathway, FAH1, FAH2 and LOH2 (LAG ONE HOMOLOG 2) led to the accumulation of sphinganine and ceramides, another class of SLs. These mutants also exhibited a PCD that was dependent on SA and was probably linked to d18:0 accumulation (König *et al*., 2021, 2012).

Sensing of pathogen-derived lipid molecules is far from being elucidated. A recent study demonstrated that the clivage product of the *Phytophthora infestans* ceramide D by the neutral ceramidase 2 is recognized by the PRR RESISTANT TO DFPM-INHIBITION OF ABSCISSIC ACID SIGNALING 2 (RDA2) (Kato *et al*., 2022). Some studies suggest that lipidic elicitors can directly interact with lipids in the plant PMs (Cordelier *et al*., 2022). For instance, ergosterol, the main sterol of fungi, is described as a general elicitor of plant defenses (Klemptner *et al*., 2014). Due to its lipidic nature and its capacity to form rafts, it is thought to interact with the PM. In *Beta vulgaris,* it can inhibit its H^+^-ATPase activity, supposedly by tempering with the NADPH oxidase present in the plant microdomains (Hao *et al*., 2014; Rossard *et al*., 2010).

While there are abundant reports on the links between LCBs and PCD (Ali *et al*., 2018; Huby *et al*, 2020), only few data on the use of exogenously applied LCBs and their mode of action are available (Glenz *et al*., 2019; Gutiérrez-Nájera *et al*., 2020; Magnin-Robert *et al*., 2015; Peer *et al*., 2011; Saucedo-García *et al*., 2011, 2023; Shi *et al*., 2007). In previous studies, we and others have shown that the co-infiltration of *Arabidopsis* leaves with an LCB and *Pst AvrRpm1* does not induce an HR (Glenz *et al*., 2022; Magnin-Robert *et al*., 2015). In order to decipher the underlying mechanism, the activation of canonical defense responses and antibacterial effect of the LCB were analyzed. Moreover, we showed similar phenotype with other effectors that share the myristoylation sequence with AvrRpm1. Finally, biophysical analyses demonstrated that sphinganine can interact with the plant biomimetic PM and disturb its organization as well as its mechanical features.

## RESULTS

### Effect of *Pst AvrRpm1* and sphinganine co-infiltration on *Arabidopsis* cell death

We and others previously demonstrated that the co-infiltration of *Pst AvrRpm1* and d18:0 or t18:0, respectively, does not induce an HR in contrast to what happens with an infiltration of an avirulent bacteria or LCB alone (Magnin-Robert *et al*., 2015; Glenz *et al*., 2022). Therefore, *Pst* expressing effectors other than AvrRpm1 were selected to elucidate whether the absence of visible HR was dependent on the effector. AvrRpt2 was chosen because, like AvrRpm1, it is associated to the PM with RIN4 (Boyes *et al*., 1998; Mackey *et al*., 2003; Xin *et al*., 2018). AvrRps4 was also tested because of the cytoplasmic localization of its target, RPS4 (Gassmann *et al*., 1999; Wirthmueller *et al*., 2007).

As expected, treatment with *Pst* DC3000 co-infiltrated or not with d18:0 triggered typical disease symptoms and infection with *Pst AvrRpm1*, *Pst AvrRpt2* or *Pst AvrRps4* alone, gave rise to a typical HR. Whereas co-infiltration of *Pst AvrRpm1* and d18:0 did not induce a visible HR as expected, the other tested bacteria displayed an HR when co-infiltrated with the sphingolipid (Fig. 1A). Changes in conductivity were then used as a marker of plant cell death, as PM damages result in strong electrolyte leakage (Kawasaki *et al*., 2005) (Fig. 1B). All treatments triggered an increase in conductivity between 4 and 24 hpi, except for co-infiltration with *Pst AvrRpm1* and d18:0, displaying a conductivity stable over time and similar to the control plants (Fig. 1B). These findings are consistent with our previous study (Magnin-Robert *et al*., 2015) and corroborated with phenotypic observations in Fig. 1A.

**Figure 1.**
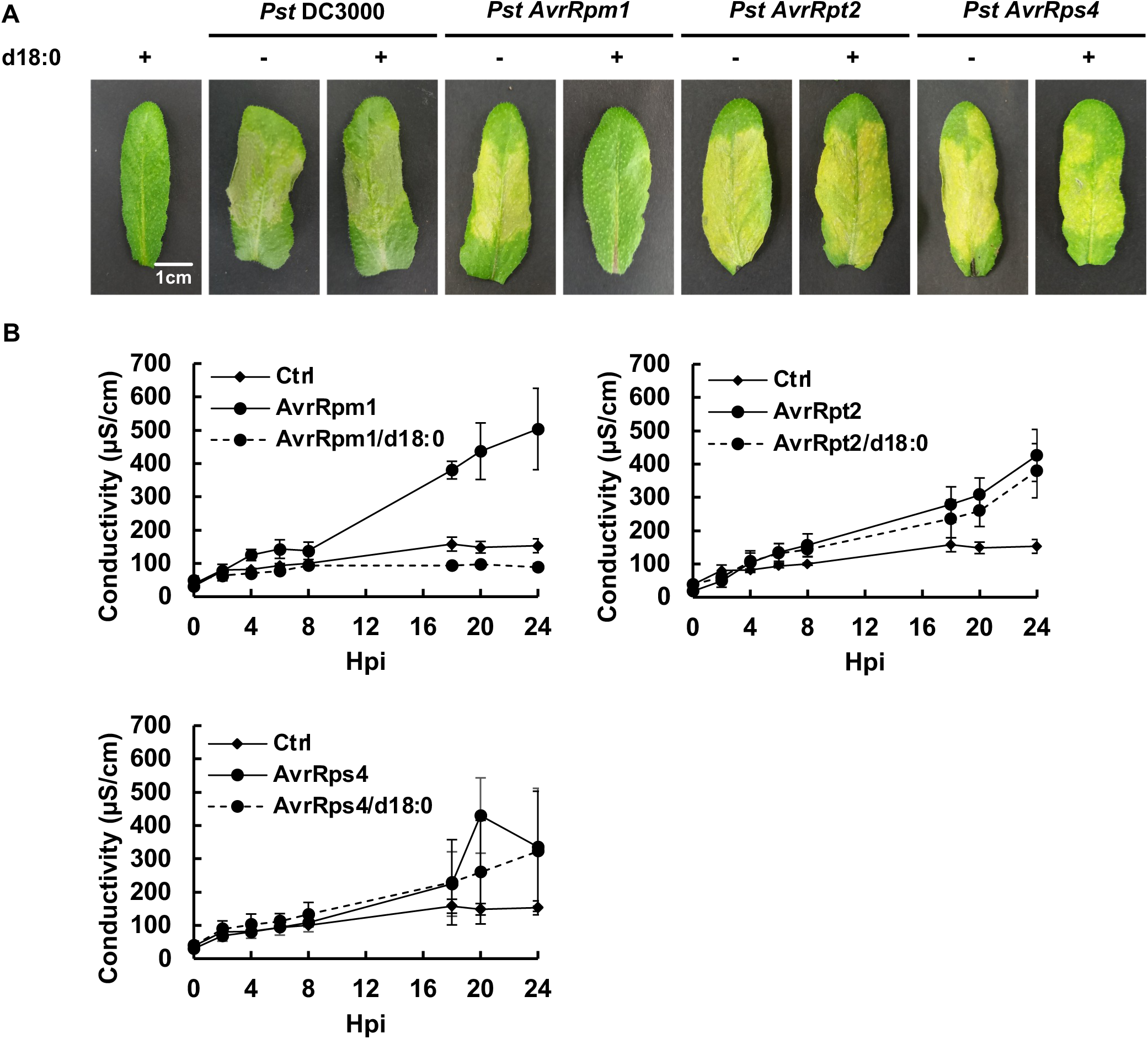
Effect of co-infiltration of *Pst* and d18:0 on cell death. (A) Representative leaves of Col-0 WT 48 hours after infiltration with either *Pst* DC3000, *Pst AvrRpm1*, *Pst AvrRpt2*, *Pst AvrRps4* alone or co-infiltrated with d18:0. The experiment was repeated at least 5 times with similar results (n=5). (B) Ion leakage measurements over 24 hours following infiltration with *Pst AvrRpm1*, *Pst AvrRpt2* or *Pst AvrRps4*, alone or with d18:0. Data are reported as means ± SD (n=3). The experiment was repeated three times with similar results.

### *In vitro* and *in planta* growth of *Pst AvrRpm1* in presence of sphinganine

As a potential deleterious effect of d18:0 on the bacteria could not be excluded, *in vitro* bacterial growth measurements were undertaken with or without the sphingolipid in the culture medium (Fig. 2A). Ethanol had no effect on bacterial growth of all tested strains (data not shown). Interestingly, d18:0 had no antibacterial effect on *Pst AvrRpm1* and out of all the bacteria tested, d18:0 slightly reduced the growth rate of *Pst AvrRpt2*. Surprisingly, this bacterium does provoke cell death when co-infiltrated into leaves.

**Figure 2.**
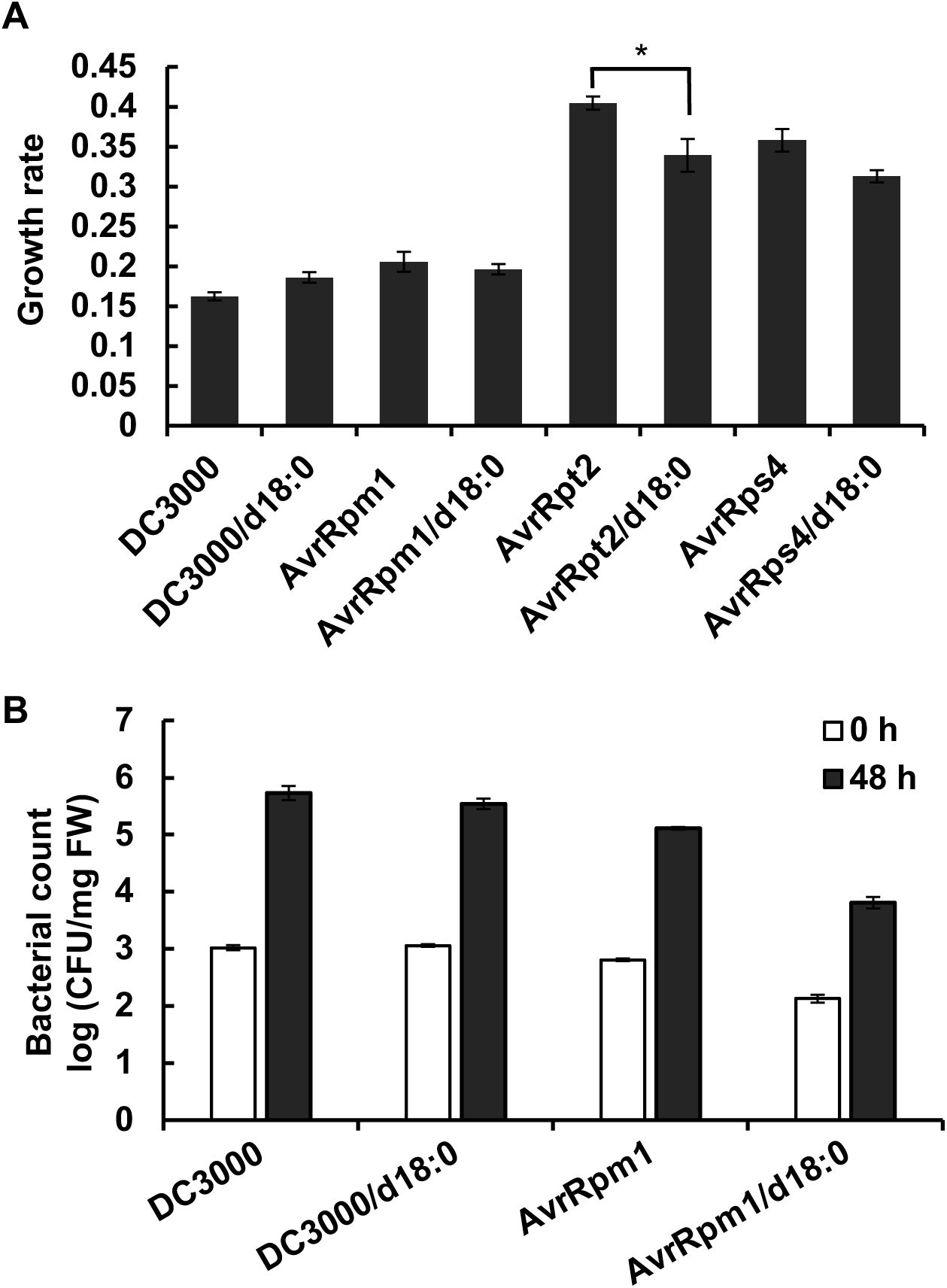
*In vitro* and *in planta* bacterial growth in presence of sphinganine. (A) Growth rate was estimated from bacterial growth curve data (OD_600_) over 48 hours, ±SD in King’s B medium. Asterisks indicate significant differences between each bacterium co-infiltrated with d18:0 or not (Wilcoxon-Mann-Whitney, *, p<0.05). (B) Bacterial growth ±SD (n = 3) at 0 and 48 hpi in *Arabidopsis* WT Col-0 leaves infected by bacteria alone or with d18:0.

To determine the effect of the co-infiltration on bacterial growth *in planta*, bacterial populations were quantified at 0 and 48 hpi (Fig. 2B). The growth of the avirulent strain *Pst AvrRpm1* was slightly reduced compared to the virulent strain, *Pst* DC3000 at 48 hpi. Furthermore, while the co-infiltration had no consequence on the development of *Pst* DC3000, it slightly reduced *Pst AvrRpm1* growth at 48 hpi. This bacterial count was however two-fold higher at 48 hpi than at 0 hpi, like in the other conditions (Fig. 2B). Collectively, these data suggest that d18:0 does not have an antibacterial effect that could explain the absence of HR when *Pst AvrRpm1* is co-infiltrated with d18:0. Moreover, while *Pst AvrRpm1* continues its development inside the plant, it is however incapable of triggering an HR.

### Extracellular ROS production by *Arabidopsis* in response to infection by *Pst*

Strong and sustained extracellular ROS production is an early defense event generally linked to HR. We therefore monitored their production by *Arabidopsis* in response to *Pst AvrRpm1*, *Pst AvrRpt2* and *Pst AvrRps4* (Fig. 3). No significant difference was found between ROS production triggered by bacteria alone or mixed with ethanol (data not shown). As already observed by Magnin-Robert *et al*. (2015), co-treatment of *Pst AvrRpm1* but also of *Pst AvrRpt2* and *Pst AvrRps4* with d18:0 reduced extracellular ROS production by approximatively 6-, 2- and 1.7-fold, respectively (Fig. 3). Only the co-infiltration of *Pst AvrRpm1* and d18:0 did not result in an HR (Fig. 1) but ROS levels appear to be dramatically affected even with other bacteria like *Pst AvrRps4*. These results suggest that reduced ROS production is not directly linked to the absence of HR.

**Figure 3.**
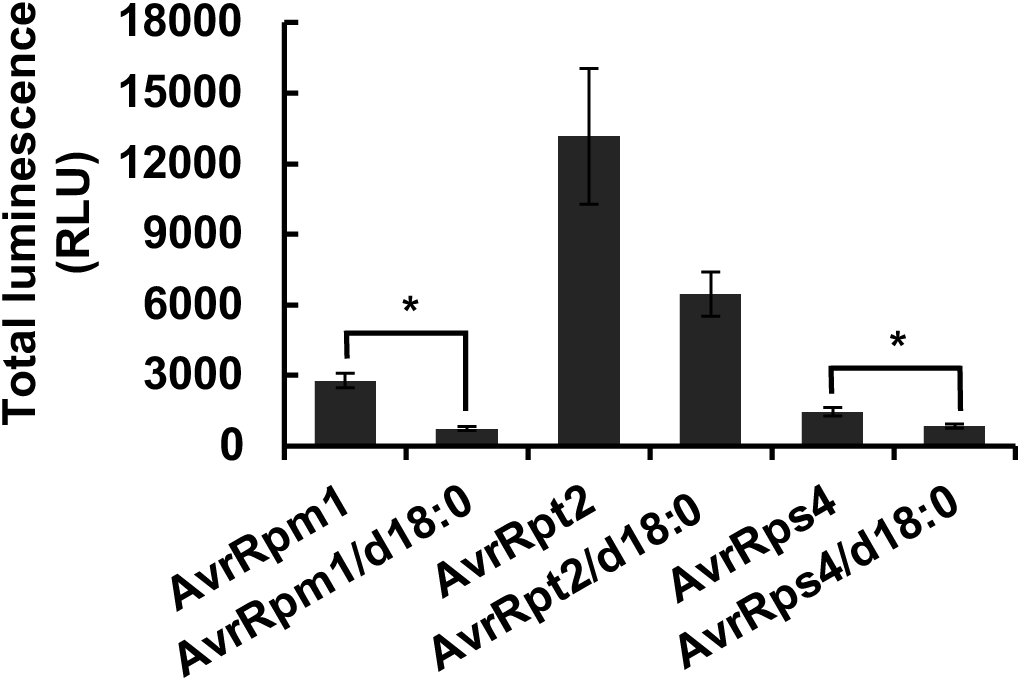
Extracellular ROS production in response to bacterial infection. Extracellular ROS production measured in presence of *Pst AvrRpm1*, *Pst AvrRpt2 or Pst AvrRps4* with or without d18:0 in the medium. Data represent the sum of RLU (Relative Light Units) ± SEM (n = 6) over 90 min. The experiment was performed three times with similar results. Asterisks indicate significant differences between each bacterium co-infiltrated with d18:0 or not (Wilcoxon-Mann-Whitney; *, p<0.05).

### Effects of *Pst AvrRpm1* and sphinganine co-infiltration on *Arabidopsis* late defense responses

Considering the effects of d18:0 on ROS production and cell death development, other markers of plant defense were studied in response to co-infiltration. The accumulation of the phytoalexin camalexin as well as the hormones SA and JA-Ile was followed. The expression of *PATHOGENESIS-RELATED GENE 1* (*PR1*) and *VEGETATIVE STORAGE PROTEIN 1* (*VSP1*), marker genes of SA and JA-Ile signaling pathways, respectively, was also analyzed. Control infiltration did not induce camalexin production (Fig. 4A). At 48 hpi, *Pst* DC3000 inoculation with or without d18:0 triggered similar camalexin accumulation. However, d18:0 co-infiltration significantly reduced camalexin production in response to *Pst AvrRpm1* since it decreased from ∼250 nmol/g FW to a similar level observed in controls (∼3 nmol/g FW) at 48 hpi (Fig. 4A). Overexpression of *PR1* was observed 48h after infiltration with each *Pst* strain alone (Fig. 4C), which correlated with the accumulation of SA (Fig. 4E). Neither SA accumulation nor *PR1* expression was induced at earlier time points. The use of an *Arabidopsis* reporter line *PR1::GUS* confirmed the induction of *PR1* expression in treated leaves (Fig. 4B). Both SA accumulation and *PR1* overexpression are repressed when leaves are co-infiltrated with d18:0, leading to similar levels as in control plants (Fig. 4, C and E). *VSP1* is weakly expressed in response to *Pst* DC3000 but overexpressed when this bacterium is co-infiltrated with d18:0 (∼ 9-fold and ∼ 6-fold more at 6 and 48 hpi, respectively) (Fig. 4D). JA-Ile levels are however higher with the bacterium alone than co-infiltrated (Fig. 4F). When *Arabidopsis* was inoculated with *Pst AvrRpm1*, *VSP1* was slightly overexpressed at 48 hpi but repressed when co-infiltrated with d18:0 (Fig. 4D). JA-Ile levels were concomitantly low (Fig. 4F), as expected from plants challenged with avirulent *P. syringae* (Mur *et al*., 2006). Overall, co-infiltration of *Pst AvrRpm1* with d18:0 significantly lowers the amount of camalexin and both defense hormone signaling pathways.

**Figure 4.**
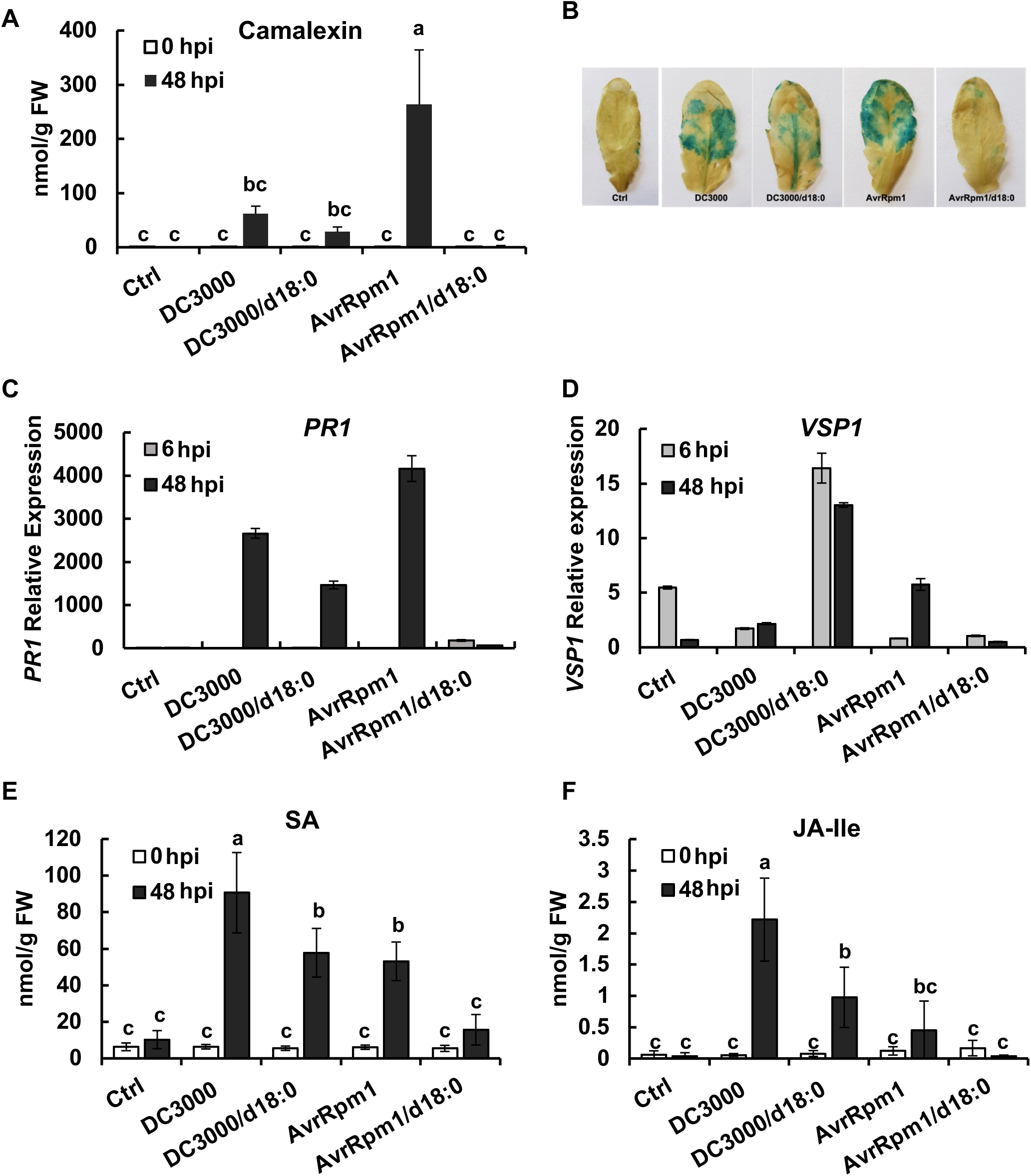
Late plant defense events in response to bacterial infiltration. *Pst* DC3000 or *Pst AvrRpm1* were co-infiltrated with or without d18:0. (A) Camalexin was quantified at 0 and 48 hpi (FW: fresh weight). Different letters indicate significant differences (ANOVA, p<0.05, one way, n=5). (B) Representative leaves of *PR1::GUS* lines 48 hpi. (C) and (D) Relative expression of *PR1* and *VSP1*, respectively, followed 6 and 48 hpi. Results are expressed as the fold increase in transcript level compared with the untreated control (0h), referred to as the x1 expression level. Experiments were repeated 3 times with similar results. (E) and (F) SA and JA-Isoleucine (JA-Ile) were quantified at 0 and 48 hpi. Different letters indicate significant differences (ANOVA, p<0.05, one way, n = 5).

#### Involvement of RPM1 and RIN4 in the absence of HR

Once injected into the cytoplasm of the host, AvrRpm1 triggers the phosphorylation of RIN4, inducing the activation of RPM1 and subsequently the ETI leading to an HR (Nimchuck *et al*., 2000). To elucidate whether RPM1 and / or RIN4 were involved when the bacterium is co-infiltrated with d18:0, inoculations of *Pst AvrRpm1* were performed on *Arabidopsis* mutants *rpm1-3,* and *rpm1-rps2-rin4,* as well as the complemented line *RPM1-myc rpm1*. Infiltration of the bacteria alone induced cell death and co-infiltration of the complemented line displayed no symptoms, as expected (Fig. 5A). Despite lacking RPM1 and / or RIN4, all mutants showed the same phenotype as the WT when co-infiltrated with d18:0 as they did not exhibit visible cell death (Fig. 5A). It has been previously demonstrated that RPM1 is degraded during the first hpi, coincidently with the onset of the HR (Boyes *et al*., 1998). To further elucidate the role of RPM1 and RIN4, *RPM1-myc rpm1* plant was used to monitor the degradation of RPM1 following inoculation (Fig. 5B). Western blot analysis showed that while RPM1 was degraded from 4 to 6 hpi when *Pst AvrRpm1* was infiltrated alone, the protein was still detected 24 hpi when the bacteria was co-infiltrated (Fig. 5B). Altogether, these data suggest that the mechanism underlying the absence of HR occurs upstream of RPM1 degradation.

**Figure 5.**
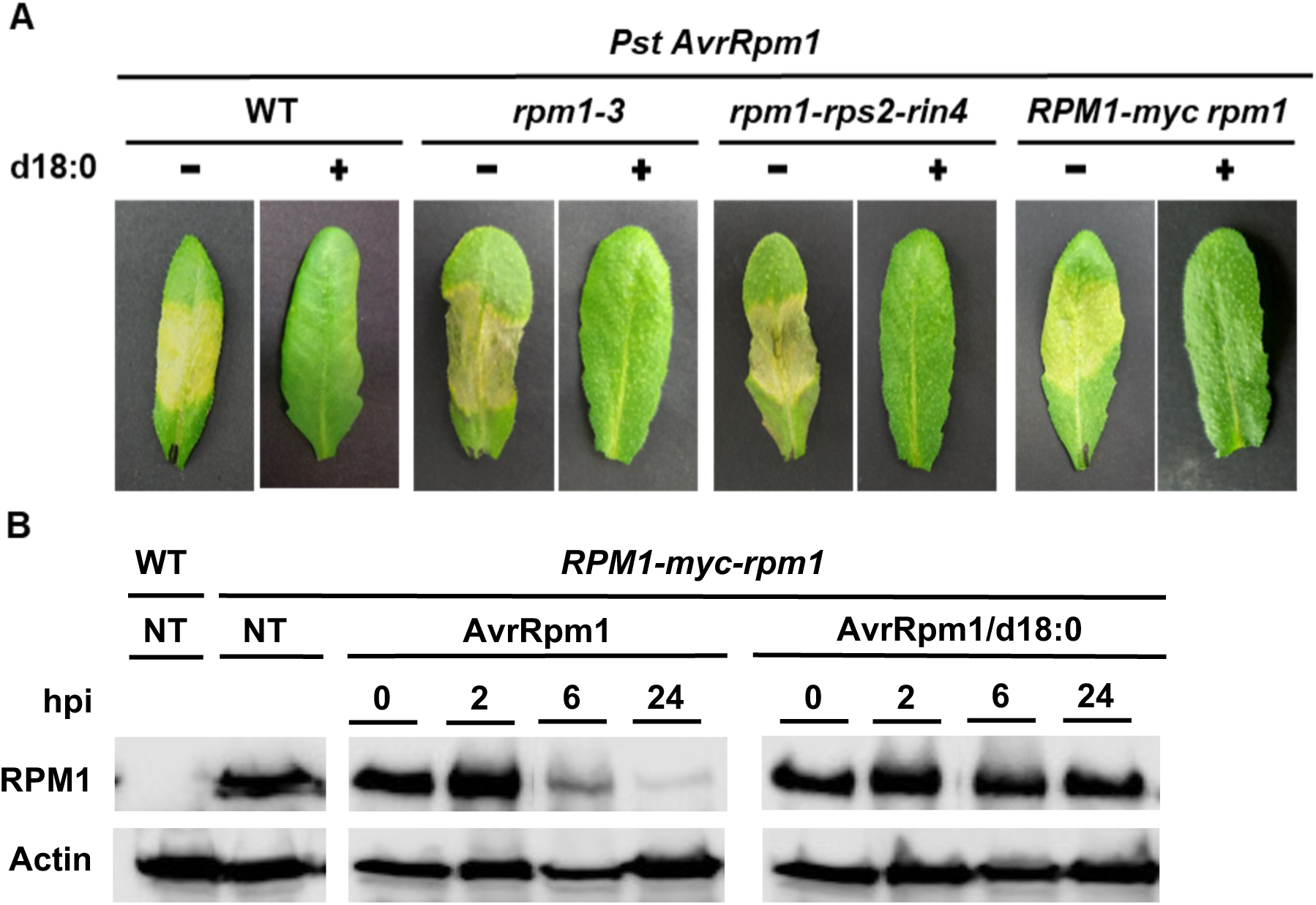
Involvement of RPM1 in the absence of defense responses. (A) Photographs represent symptoms observed 72 hpi in WT, *rpm1-3*, *rpm1-rps2-rin4* and *RPM1-myc rpm1* lines treated with or without d18:0. (B) Western blot of RPM1 protein in WT and *RPM1-myc rpm1* line at 2, 6 or 24 hpi. Normalization was carried out with Actin. NT refers to non-treated samples. All experiments were repeated at least 3 times with similar results.

### Effect of *Pst AvrB* and *Pst AvrPphB* co-infiltration with sphinganine on *Arabidopsis* defense response

The three effectors AvrRpm1, AvrB and AvrPphB share an eukaryotic *N*-myristoylation site in their protein sequences that allows them to be redirected to the PM once they have entered their host (Nimchuk *et al*., 2000). Like AvrRpm1, AvrB interacts with RPM1 and RIN4 (Grant *et al*., 1995; Lee *et al*., 2004), and AvrPphB with RPS5 and PBS1, both located at the PM (Ade *et al*., 2007; Pottinger and Innes, 2020). As already demonstrated (Warren *et al*., 1998; Whalen *et al*., 1991), infiltration of *Pst AvrB* or *Pst AvrPphB* triggers an HR. No cell death was visible when both *Pst AvrB* and *Pst AvrPphB* were co-infiltrated with d18:0, suggesting HR was not triggered in treated plants, similarly to *Pst AvrRpm1* (Fig. 6A). Further tests were conducted to determine if d18:0 had an *in vitro* effect on these bacteria. Their growth rate remained unchanged after d18:0 addition (Fig. 6B), confirming that d18:0 has no antibacterial effect on these two bacteria. Extracellular ROS production by *Arabidopsis* in response to *Pst AvrB* and *Pst AvrPphB* was also monitored (Fig. 6C). As for *Pst AvrRpm1*, d18:0 significantly reduced this production in response to both bacteria (∼ 2-fold for *Pst AvrB* and ∼ 3-fold for *Pst AvrPphB*).

**Figure 6.**
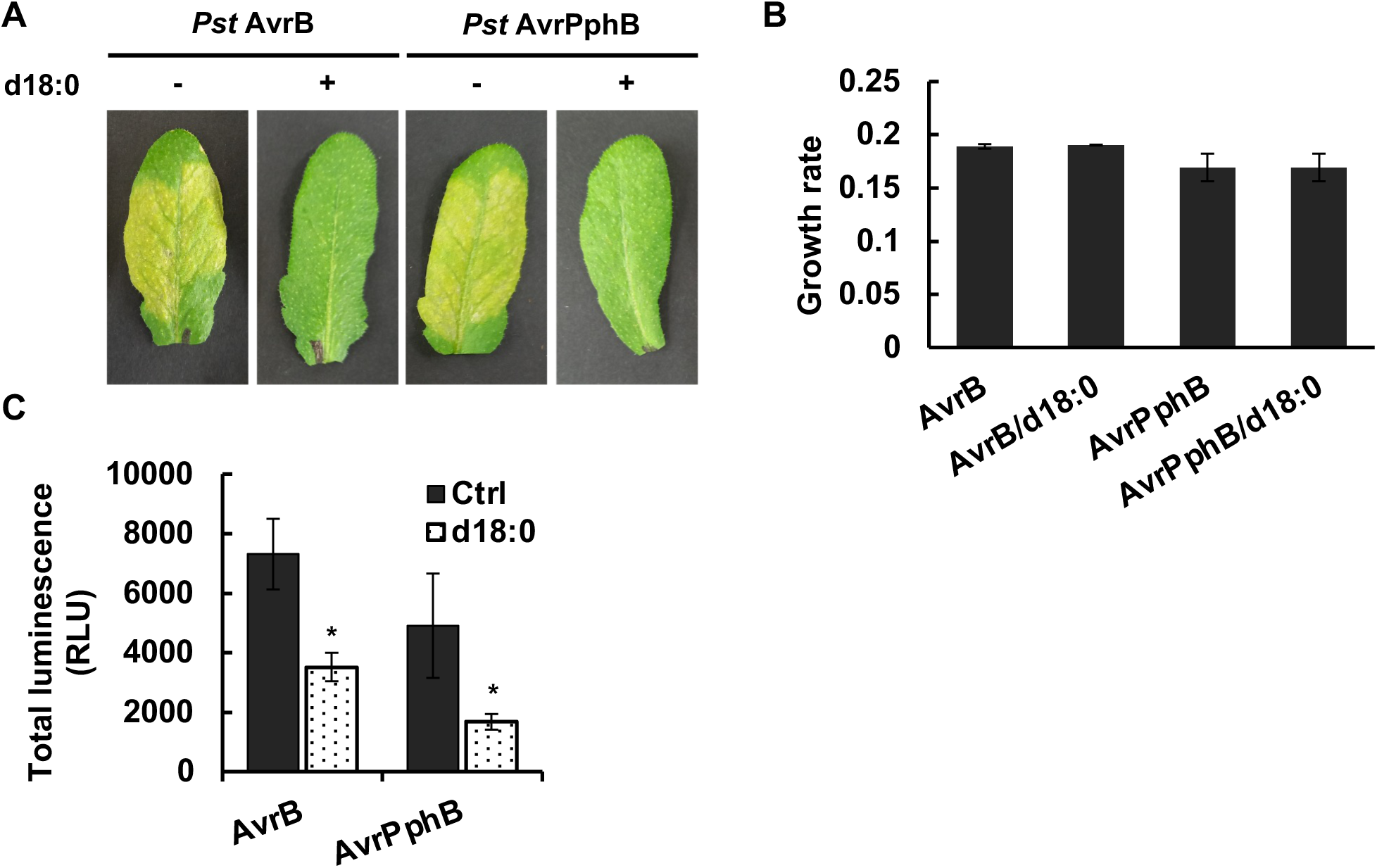
Effect of *Pst AvrB* and *Pst AvrPphB* co-infiltration with d18:0 on plant defense responses. (A) Photographs represent symptoms observed 48 hours after WT plant infiltration with either *Pst AvrB* or *Pst AvrPphB* alone or with d18:0. The experiment was repeated at least 3 times with similar results (n = 3). (B) Growth rate estimated from bacterial growth curve data (OD_600_) over 48 hours, ±SD in King’s B medium. (C) ROS production measurement in presence of *Pst AvrB* or *AvrPphB* with or without d18:0. Data represent the sum of RLU (Relative Light Units) ± SEM (n = 6) over 90 min. Experiments were performed three times with similar results. Asterisks indicate statistical differences (Wilcoxon-Mann-Whitney; *, p<0.05).

### *NMT1* relative expression following *Pst AvrRpm1* and sphinganine co-infiltration

In *Arabidopsis*, two enzymes, NMT1 and NMT2, are responsible for *N*-myristoylation (Pierre *et al*., 2007). As the corresponding mutants displaying severe phenotypes, they were thus not suitable for our experiments. We used qRT-PCR to determine expression patterns of *NMT1* and *NMT2* in response to infiltration of *Arabidopsis* WT leaves with *Pst* DC3000 or *Pst AvrRpm1*, either alone or with d18:0 (Fig. 7). Unfortunately, the expression of *NMT2* was too low to be detectable. At 6 hpi, *NMT1* expression in plants infiltrated with *Pst* DC3000 alone or with d18:0 was similar to that of control samples but was induced at 48 hpi (∼ 2-fold). When plants were infiltrated with *Pst AvrRpm1* alone, a stimulation of *NMT1* expression is observed during the time-course experiment to reach a ∼2.2-fold induction at 48 hpi (Fig. 7). However, when plants were co-infiltrated with *Pst AvrRpm1* and d18:0, no induction could be observed during the time-course experiment (Fig. 7). Thus, a decrease of ∼1.7-fold is observed 48 hpi between *Pst AvrRpm1* infiltrated alone and with d18:0. Overall, these results suggest that co-infiltration with d18:0 modifies *NMT1* expression in response to infection with *Pst AvrRpm1*.

**Figure 7.**
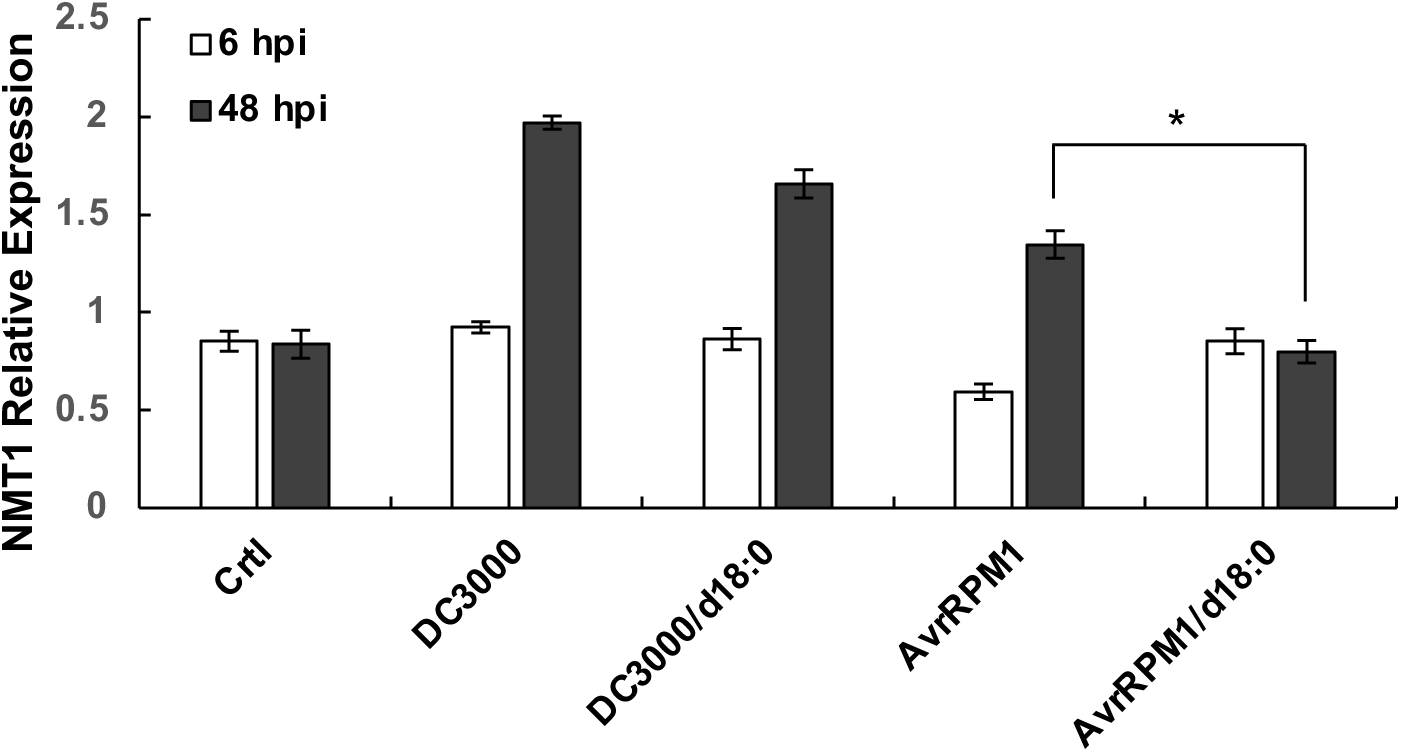
*NMT1* expression in response to co-infiltration of bacteria and sphinganine. Relative expression of *NMT1*, (N*-myristoyltransferase 1*) was followed at 6 and 48 hpi of *Pst* DC3000 or *Pst AvrRpm1* with or without d18:0. Results are expressed as the fold increase in transcript level compared with the untreated control (0h), referred to as the x1 expression level. Experiments were repeated three times with similar results. Asterisks indicate significant differences between each bacterium co-infiltrated with d18:0 or not (Wilcoxon-Mann-Whitney; *, p<0.05).

### Interaction capacity of sphinganine with plant plasma membrane-mimicking models

It has been shown that LCB sphingosine (d18:1) can increase the permeability of liposomal and erythrocyte ghost membranes to aqueous solutes (Contreras *et al*., 2006). Based on this finding, d18:0 was hypothesized to interact with the lipids of plant PM. To investigate this hypothesis, we conducted *in silico* and *in vitro* biophysical analyses using membrane plant models. Thanks to their defined structure and controlled lipid composition, these models were found to be useful in understanding how external molecules affect the organization and mechanical properties of the membranes (Deleu *et al*., 2014; Furlan *et al*., 2020; Cordelier *et al*., 2022).

The Impala modeling method was used to predict the ability of d18:0 to insert into an implicit model lipid bilayer and to determine the preferable location and positioning of d18:0 within a membrane bilayer (Fig. 8). The more negative the energy is, the more favorable the insertion will be. A sharp drop of energy was observed as d18:0 moved from the aqueous phase to the membrane surface (up to ∼-10 Kcal/mol), indicating that it could easily insert into membranes. The energy was close to 0 Kcal/mol in the hydrophobic region of the membrane, suggesting that d18:0 has a less favorable interaction with this region compared with the lipid bilayer hydrophilic region (Fig. 8A). Nevertheless, the non-positive energy value in the hydrophobic region of the bilayer suggests that d18:0 could flip-flop between the two bilayer leaflets. A large increase in energy (from ∼-10 Kcal/mol to +6.7 Kcal/mol) is also observed when d18:0 moves from the inner bilayer leaflet to the aqueous medium, which strongly suggests that d18:0 has no propensity to cross the membrane and enter into the cytoplasm. The preferred positioning of d18:0 within the membrane is shown in Fig. 8B. Its lipid chain lies within the hydrophobic part of the membrane while the amine and alcohol groups are located in the hydrophilic part of the bilayer. Altogether, this indicates that d18:0 can interact with membranes and remains there. To support these simulation results, experimental analyses were performed using large unilamellar vesicles (LUVs) that mimic plant PM, consisting of palmitoyl-2-linoleoyl-*sn*-glycero-3-phosphocholine (PLPC), glucosylceramide (GluCer) and β-sitosterol (sito) as the plant PM model phospholipid, sphingolipid and sterol respectively (Deleu *et al*., 2014; Rondelli *et al*., 2021; Deboever *et al*., 2022).

**Figure 8.**
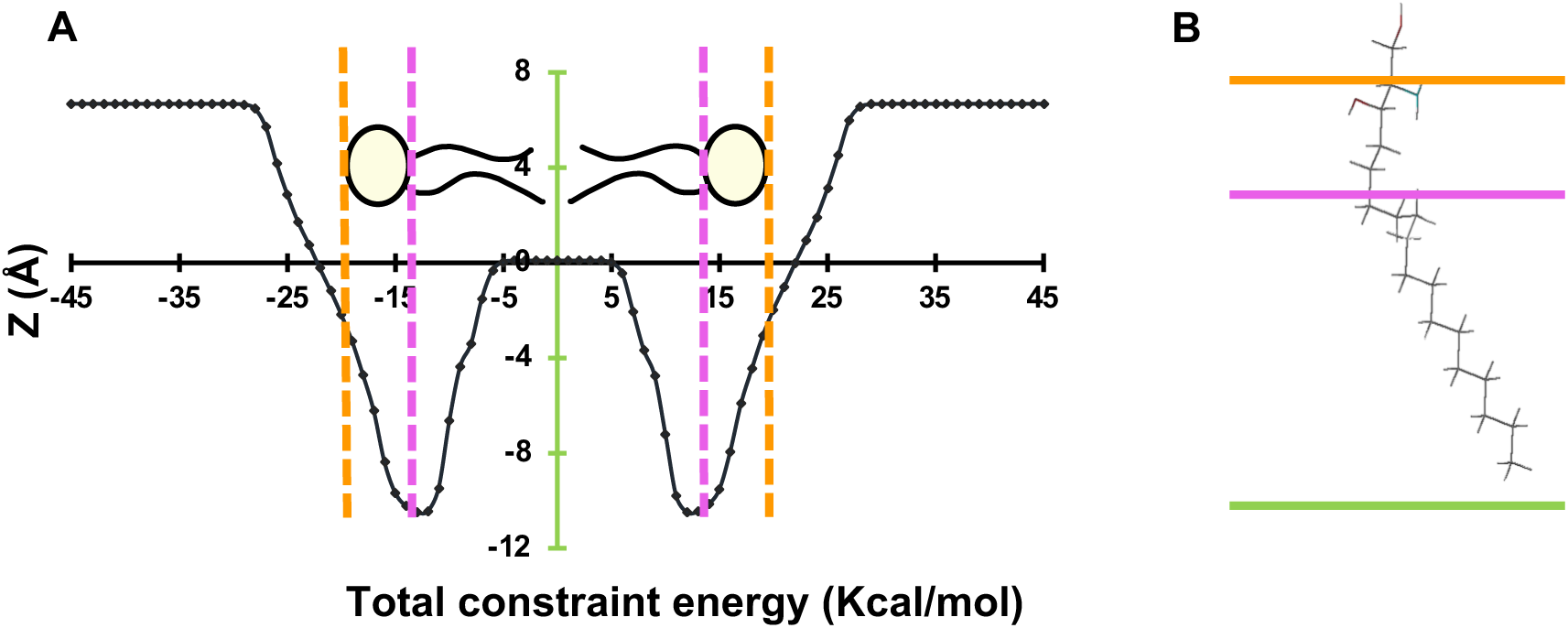
Interaction of sphinganine with an implicit modeled bilayer. (A) The Impala simulation calculates the energy restraints of a lipid bilayer as d18:0 goes through it. The X-axis corresponds to the position of the center of mass along the Z-axis. From left to right: the interface between the bilayer and the aqueous phase (orange), the interface between the polar head and the alkyl chain (pink) and the center of the bilayer (green). (B) The most energetically favorable orientation of d18:0 in the modeled bilayer. The lines represent the same planar surfaces as in A.

Results from ITC experiments showed a negative and decreasing heat flow after each LUV injection into the d18:0 solution (Fig. 9A), indicating that d18:0 interacts with plant PM-mimicking LUVs. The thermodynamic parameters calculated from the fitting of raw data are presented in Fig. 9B. The binding coefficient K was relatively high (>80 mM^-1^) suggesting that d18:0 has a strong affinity for plant-mimicking lipid membranes confirming the low probability that it emerges in the intracellular medium. This affinity for plant PM-like liposomes is higher than that observed for other plant-eliciting molecules such as linoleic and linolenic acid hydroperoxides (<3 mM^-1^; Deleu *et al*., 2019), and rhamnolipid bolaforms (7.5 mM^-1^; Luzuriaga-Loaiza *et al*., 2018). The binding reaction of d18:0 to LUVs was spontaneous as the molar free energy ΔG was negative, exothermic as the molar enthalpy change ΔH was negative and generated a positive change of the molar entropy (TΔS>0). The higher absolute value of the molar entropy (TΔS>0) compared to the absolute value of the molar enthalpy change (ΔH), indicates that the interactions is entropy-driven, i.e. mostly of hydrophobic nature. Altogether, these results suggest that d18:0 is attracted by and can interact with plant lipid membranes.

**Figure 9.**
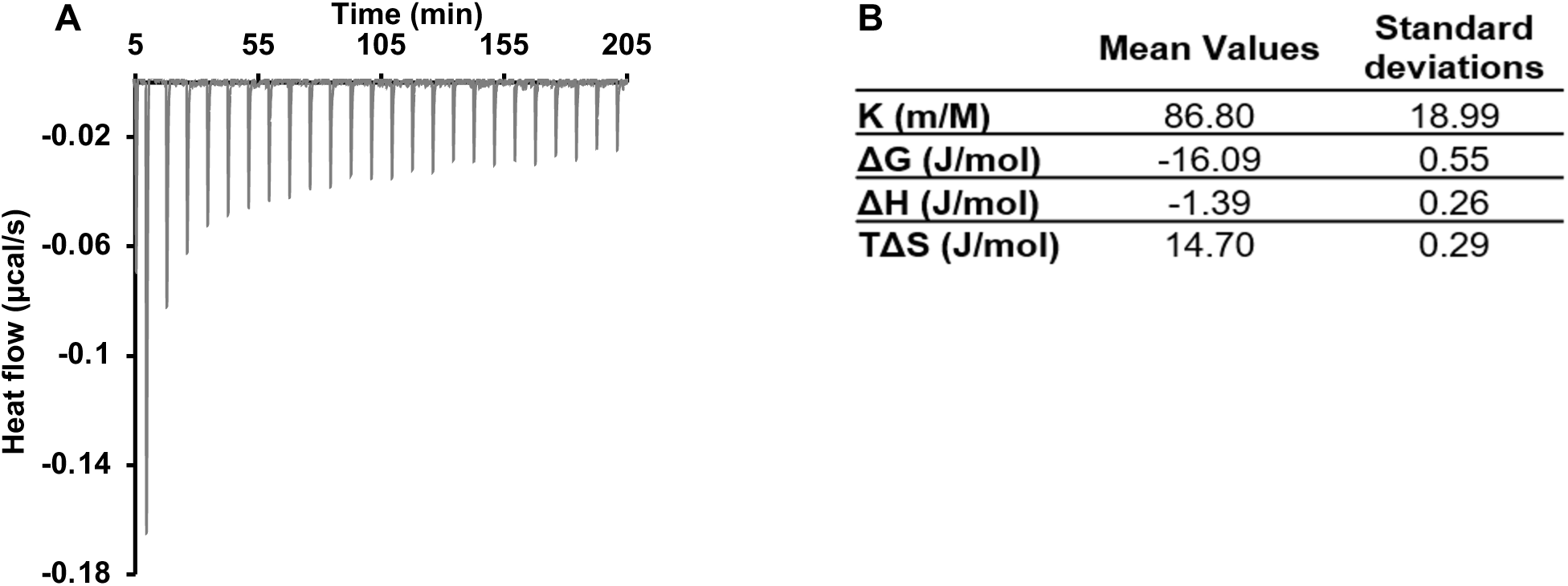
Interaction of sphinganine with plant-like liposomes. (A) Raw data from ITC experiments at 26°C. Each peak corresponds to a single injection of a 1 mM PLPC, glucosylceramide and sitosterol (60:20:20, molar ratio) LUV suspension into a solution containing 20 µM of d18:0. (B) Calculated thermodynamic parameters for this interaction. These values are means of two independent experiments. K represents the binding coefficient (the affinity of d18:0 for the LUVs), ΔG represents the molar free energy, ΔH, the molar enthalpy change, and TΔS, the molar entropy change.

### Lipid specificity of the interaction between sphinganine and plant plasma membrane-mimicking models

To investigate specific interactions between d18:0 and lipids representative of plant PM, the Langmuir trough technique was used to analyze the adsorption capacity of d18:0 into monolayers of each individual lipid. In this technique, a lipid monolayer is initially formed at an air-water interface, and then d18:0 is injected into the subphase beneath the monolayer. The increase in surface pressure (ΔП) at a constant trough area was monitored to determine the adsorption of d18:0 into the lipid monolayer at different initial surface pressures (П_i_) (Supplemental Fig. S1). Two parameters were determined from these data (Fig. 10A): (i) the differential surface pressure (dП_0_), which provides information on the attractive effect of the lipids on d18:0, and (ii) the maximal insertion pressure (MIP), which reflects the ability of d18:0 to penetrate into the lipid monolayer (Deleu *et al*., 2014). Upon insertion of d18:0 onto monolayers formed with individual plant PM lipids, dП_0_ values were all positive and similar for all compositions, indicating that all tested lipids exert a similar attraction on d18:0 (Fig. 10A). MIP values were all higher than the lateral pressure supposed to prevail into biological membranes (30 – 35 mN/m) (Marsh, 1996), meaning that d18:0 may be able to adsorb into natural plant membranes.

**Figure 10.**
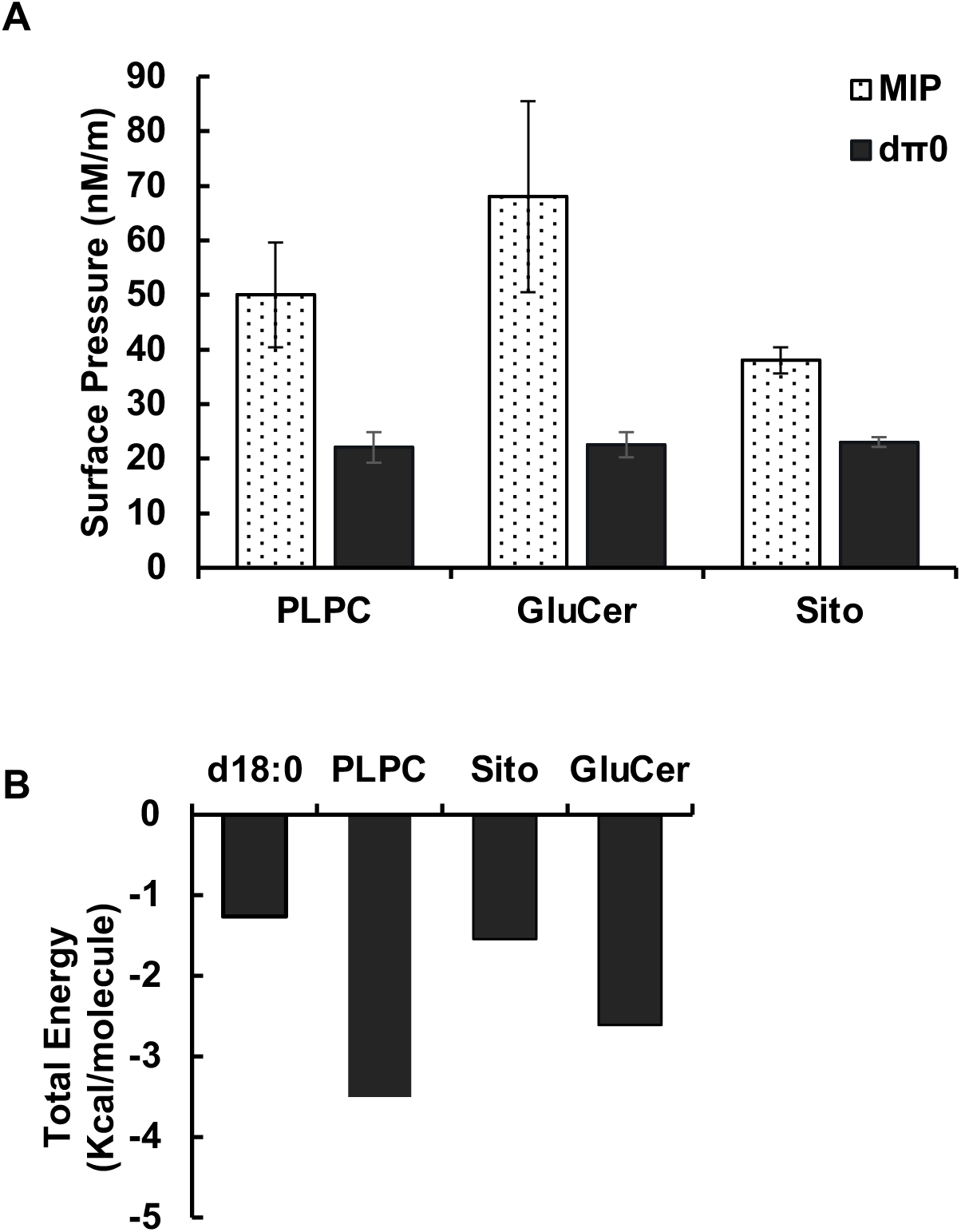
Specificity of sphinganine interaction with various lipids representative of the plant plasma membrane. (A) Adsorption of d18:0 into monolayers of plant PM lipids: PLPC, glucosylceramide (GluCer) and β-sitosterol (Sito). d18:0 was injected at 1.25 µM in a subphase (Tris HCl, 10 mM, pH 7.4), underneath the lipid monolayer. The maximal insertion pressure (MIP) reflects the penetration power of d18:0 into the lipid monolayer and the differential П (dП0) indicates the attracting effect the lipids have on it. (B) Total energies of interaction of d18:0 alone or with lipids representative of the plant plasma membrane were calculated with the Hypermatrix docking method. Lipids used were PLPC, GluCer, and Sito.

Lipid specificity of the interaction was also studied by an *in silico* approach using the Hypermatrix docking method. This method calculates the energy of interaction between d18:0 and individual plant lipids at a hydrophilic/hydrophobic interface, indicating whether this interaction is energetically favorable or not (Brasseur *et al*., 1987; Deleu *et al*., 2014). Results showed that the interaction between d18:0 and all plant lipids was favorable as all interaction energies are negative (Fig. 10B). Interestingly, the interaction energies between d18:0 and plant lipids were more favorable than those observed for d18:0 with itself. With this method, interaction between d18:0 and PLPC or GluCer appeared to be more favorable than with β-sitosterol.

Altogether, these data showed that d18:0 is attracted by and could interact with lipids of plants. There is however no clear trend to determine a preferential interaction of d18:0 with a particular lipid.

### Modification of the organization and mechanical properties of plant plasma membrane-mimicking models induced by sphinganine

Since d18:0 can interact with lipids found in plant PMs, additional experiments were conducted to investigate how these interactions could impact the mechanical properties and the organization of plant PMs.

Two aspects of membrane mechanics were considered: membrane fluidity and membrane elasticity. Membrane fluidity refers to the degree of lipid packing and the mobility or dynamics of lipids within the membrane, which can be characterized by the order level parameter (Gerbeau-Pissot *et al*., 2014). Membrane elasticity corresponding to the resistance of a material to deformation when subjected to an applied force was determined by measuring the Young’s modulus. The higher the Young’s modulus, the stiffer and less deformable the object is (Magazzu and Marcuello, 2023). Although membrane fluidity can be influenced by the membrane elasticity, there is no direct quantitative relationship between Young’s modulus and membrane fluidity or order level and therefore must be considered as separate aspects of membrane behavior.

To analyze the order level of *Arabidopsis* PM in response to d18:0, *Arabidopsis* leaf protoplasts were stained with the dye di-4-ANEPPDHQ. The order level of the membrane, revealed by fluorescence imaging, was quantified using the red/green ratio of the membrane (RGM) (Dinic *et al*., 2011; Gerbeau-Pissot *et al*., 2014). Observations of protoplasts indicated a significant decrease in RGM compared to controls (from 0.99 to 0.80 and 0.74 for concentrations of 2 µM and 5 µM d18:0, respectively) (Fig. 11A-D), with no statistical difference between the two concentrations of d18:0. The RGM profile of a 4 µm fragment of PM from controls appeared homogenous whereas those from sphingolipid-treated protoplasts is more heterogenous (Fig. 11E-G). These findings suggest that the order level increased in response to the LCB. This increase in membrane order level in the presence of d18:0 was confirmed by the greater variation in the generalized polarization of Laurdan on biomimetic liposomes compared to the control ethanol, and this as soon as d18:0 came into contact with the lipid bilayer (Fig. 11H). Laurdan is a fluorescent probe sensitive to membrane lateral packing, a higher generalized polarization corresponding to a greater lipid packing (Parasassi *et al*., 1991).

**Figure 11.**
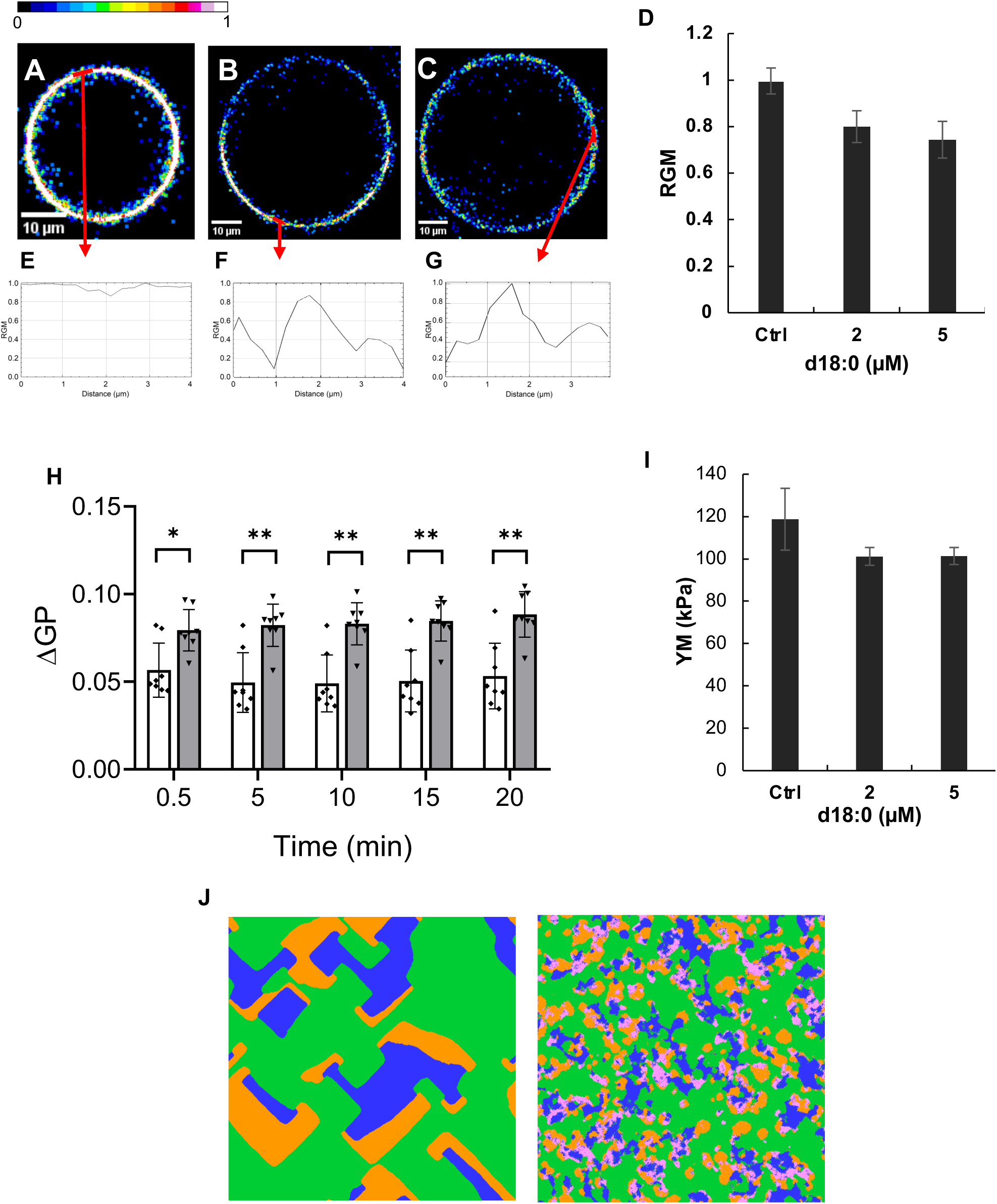
Mechanical properties and organization of the plant plasma membrane in response to sphinganine treatment. (A) to (C) Fluorescence red/green ratio images by multispectral confocal microscopy of a single-cell PM labeled with di-4-ANEPPDHQ after control (A), 2 µM d18:0 (B) or 5 µM d18:0 (C) treatment. (D) Red to green ratio of the PM (RGM) labeled with di-4-ANEPPDHQ treated with control (Ctrl), 2 µM or 5 µM d18:0. The asterisk indicates a significant difference compared to the control (P value < 0.05). (E) to (G) RGM profile on a 4 µm distance on PM from protoplasts after control (E), 2 µM d18:0 (F) or 5 µM d18:0 (G) treatment. (H) Variation of laurdan generalized polarization (ΔGP) measured in PM-mimicking liposomes (50 µM PLPC:Glucosylceramide: β-sitosterol (60:20:20 molar ratio)) at several time points after control (white bars) and 20 μM d18:0 treatment (grey bars). A positive ΔGP corresponds to a higher lipid packing within the bilayer. Mean ± SD of two replicates from four independent experiments. Asterisks indicate statistically significant differences with control, *P<0.05, **P<0.01, two-way ANOVA and Sidak’s multiple comparison test. (I) Young’s modulus measured on mesophyll protoplasts before or after 5 min incubation of d18:0 at two concentrations. Mean ± SD of at least 50 analyses from three independent experiments. (J) Monolayer simulations of lipid-d18:0 interactions performed with Big Monolayer where each pixel represents a molecule. (Left) Plant lipid model with PLPC in green, glucosylceramide (GluCer) in blue and sitosterol (sito) in orange (60:20:20, molar ratio). (Right) The same lipids after addition of d18:0 in pink (88% lipids, 12% d18:0).

Young’s modulus was measured by recording force-distance curves using AFM on leaf protoplasts. In the presence of d18:0, Young’s modulus decreased (Fig. 11I) indicating a reduction in membrane elasticity and a lower resistance to membrane deformation. Again, there is no concentration dependence.

Using the Big Monolayer simulation method, which is useful for studying the lateral distribution of molecules based on their mutual interaction energies, we observed that d18:0 preferentially interacted with the lipid domains initially formed by β-sitosterol and GluCer (Fig. 11J left), resulting in their fragmentation as shown in Fig. 11J right. Accordingly, fluorescence images of di-4-ANEPPDHQ-stained protoplasts (Fig. 11 A to C and E to G) demonstrated an increase in the number of small ordered domains in the presence of d18:0 compared to its absence.

## DISCUSSION

The roles of sphingolipids during PCD in response to biotic stress are still not fully understood (Berkey *et al*., 2012; Huby *et al*., 2020). Previous studies showed that co-infiltration of *Pst AvrRpm1* with the LCB d18:0 or t18:0 on *Arabidopsis* leaves did not trigger an HR on the treated leaves, with no evidence of cell death, indicating the conservation of plant PM integrity and absence of cell lysis following the co-infiltration (Glenz *et al*., 2022; Magnin-Robert *et al*., 2015). In the present work, we show that the levels of all plant immunity markers in response to co-infiltration, including those related to the plasma membrane, are similar to the control plants, despite the fact that the bacterium still develops inside plant tissues. We also report a similar effect with *Pst* expressing the effector AvrB or AvrPphB. AvrRpm1, AvrB and AvrPphB need to be myristoylated to be active. This process is disturbed during co-infiltration leading to an absence of plant defense response activation. Finally, interaction of d18:0 with lipids of the plant PM was demonstrated.

*In vitro* tests showed that this LCB does not have an antibacterial effect on *Pst AvrRpm1*, *Pst AvrB* and *Pst AvrPphB*, similarly to results obtained with *P. aeruginosa* (Fischer *et al*., 2012). t18:0 has been shown to inhibit *Pst* growth (Glenz *et al*., 2022). However, the authors used a smaller bacterial concentration and LB medium to perform their tests. Here, by using a higher bacterial concentration and a medium more favorable to *Pst* growth, the bacterial pressure was more important in our tests, which could explain the discrepancy with the data published by Glenz and colleagues (2022). *In planta* experiments showed that, even though *Pst AvrRpm1* is not capable of inducing an HR when co-infiltrated with d18:0, it is still present inside the plant and even continues to thrive, confirming the non-detrimental effect of d18:0 on the bacterium. In contrast, a pre-treatment with FB1, in order to promote endogenous LCB accumulation, inhibit *Pst* DC3000 but not *Pst AvrRpm1* growth (Saucedo-García *et al*., 2023). Moreover, *loh1* plants that accumulate endogenous levels of LCBs together with *PR1* enhanced expression, displays spontaneous cell death (Ternes *et al*., 2011). The localization (apoplastic *vs* cytoplasmic) of the LCB and the fact that FB1 or *loh1* induces or displays higher LCB but also LCB-P, ceramide or GluCer accumulation (Chen *et al*., 2008; Kimberlin *et al*., 2016; Saucedo-García *et al*., 2011) could explain this discrepancy. This could also suggest that interaction of d18:0 with the PM and its presence within the PM when infiltrated could be essential in the cell death induction. Previous results showed that co-treatment of d18:0 with *Pst* DC3000 or with *Pst AvrRpm1* drastically reduced extracellular ROS production by *Arabidopsis* (Magnin-Robert *et al*., 2015). Our results demonstrate that d18:0 reduces ROS production in response to bacteria that induce symptoms *in planta* but also to bacteria that do not induce symptoms (*Pst AvrRps4* and *Pst AvrRpm1, Pst AvrB, Pst AvrPphB,* respectively). This production remained unchanged when plants are treated with *Pst AvrRpt2*. The hypothesis that LCBs could impact ROS production by plants has already been put forward in several studies, although some of them are somewhat contradictory. Coursol *et al*. (2015) showed that exogenous addition of 20 µM LCBs reduced tobacco ROS production in response to cryptogein. In *Arabidopsis*, LCBs can induce both intracellular as well as extracellular ROS production (Glenz *et al*., 2019; Peer *et al*., 2011; Shi *et al*., 2007; Saucedo-Garcia *et al*., 2023). ROS production can be detected during both PTI and ETI (Torres, 2010). In *Arabidopsis,* it is mainly related to RbohD located in lipid microdomains of the plant PM (Hao *et al*., 2014; Mongrand *et al*., 2004). Several studies have suggested that RbohD-derived ROS production is not directly involved in plant cell death, but rather in the signaling pathways associated with plant defense responses (Lherminier *et al*., 2009; Torres *et al*., 2005). This means that even though d18:0 could alter ROS production, it may not have any impact on PCD and would therefore not be responsible for the absence of HR. Additionally, it has been already demonstrated that a lipidic elicitor is able to interact with lipid membrane microdomains (Rossard *et al*., 2010). These data suggest that LCBs might interact with and thus disturb the PM organization of *Arabidopsis* leading to the reduction of RbohD activity.

SA pathway is triggered in response to hemibiotrophic pathogens, such as *Pst*, whereas JA and its active form JA-Ile are involved in infection by necrotrophic pathogens (Li *et al*., 2019). In our study, *PR1* is significantly down-regulated upon co-infiltration with d18:0 and *Pst* DC3000, and even more so upon *Pst AvrRpm1,* which coincides with lower SA accumulation. SA repression is commonly associated with promotion of bacterial development and/or enhancement of a JA-dependent pathway (Brooks *et al*., 2005; Moore *et al*., 1989; Zhou *et al*., 1998). *P. syringae* produces coronatine, a phytotoxin that mimics JA-Ile and acts by inhibiting the accumulation of SA in order to promote bacterial growth (Zheng *et al*., 2012). Conversely, in *Arabidopsis*, coronatine-insensitive mutants that are resistant to *Pst* display higher levels of SA (Block *et al*., 2005). In *Arabidopsis*, the SA and JA pathways antagonize each other (Li *et al*., 2019; Mur *et al*., 2006). However, JA pathway markers are also down-regulated in response to d18:0/*Pst AvrRpm1* infiltration. Considering these results, we could conceive that co-infiltration represses or does not activate both SA and JA pathways in response to the avirulent bacterium because of a lowered bacterial virulence, a decreased susceptibility, and/or an altered perception of the pathogen.

AvrRpm1, AvrB, and AvrPphB effectors all possess a consensus N-terminal myristoylation sites which are required for full avirulence function (Nimchuk *et al*., 2000). Myristoylation process occurs inside the plant cell (Nimchuk *et al*., 2000) and is catalyzed by NMT (*N*-myristoyltransferase) enzymes (Pierre *et al*., 2007). Here, *NMT1* expression is significantly reduced at 24 and 48 hpi when *Pst* AvrRpm1 is co-infiltrated with d18:0. d18:0 co-infiltration could thus interfere with the myristoylation of these effectors. As a consequence, it prevents them from being redirected to the PM where their targets are and consequently preventing their activity in *Arabidopsis*. Recently, a mis-localization of RPM1 at the PM, due to a depletion in anionic phospholipids in the PM, inhibits its cell death activity (Saile *et al*., 2021). Since cell death is still induced, such depletion does not affect RPS5 function, which is constitutively PM localized due to the presence of myristoylation motifs in its sequence. RPS5 mediates the recognition of AvrPphB. Co-infiltration of this effector with d18:0 results in an absence of HR, thus excluding the involvement of a PM phospholipid dependent mechanism. PM localization and stability of NLRs is essential for their activity in the induction of cell death and immune responses (Saile *et al*., 2021). Effector localization into plant cell is also essential for their interaction with their targets and subsequent activity (Nimchuk *et al*., 2000). The inability for AvrRPM1 to interact with its target implies not only that all the downstream plant defense responses are not activated but also the absence of RPM1 degradation after co-infiltration. Moreover, mutation of consensus myristoylation sites of *AvrRPM1* or *AvrB* leads to a highly reduction of HR events and an unhindered pathogen growth (Nimchuk *et al*., 2000), consistent with our results.

All the effects of d18:0 previously mentioned, and its amphiphilic nature suggest that the molecule can interact with the plant PM. It was thus hypothesized that it could insert into the lipid fraction of the PM, as already suggested for other molecules such as fatty acid hydroperoxides, rhamnolipids, lipopeptides that interact with lipids of plant PMs (Cordelier *et al*., 2022; Deboever *et al*., 2022; Deleu *et al*., 2013, 2008; Henry *et al*., 2011; Nasir *et al*., 2010; Schellenberger *et al*., 2021). Here, biophysical studies have shown that d18:0 was attracted by, interacted with and stayed in biomimetic plant PMs. These findings were further supported by *in silico* and *in vitro* analyses as well as microscopic observations that notably showed d18:0 could disturb the lateral organization and mechanical properties of plant PM and more specifically of its microdomains. While it was suggested that exogenous LCB could cross the PM by passive diffusion (Glenz *et al*., 2019), our results suggested that d18:0 has no propensity to go into the intracellular medium. It actually possesses a flip-flop potential within the PM meaning that exogenous addition of d18:0 could disturb the PM. As previously stated, this interaction could eventually disturb defense-related proteins located within its domains (Gronnier *et al*., 2018, 2016; Huby *et al*., 2020; Mamode Cassim *et al*., 2019; Nagano *et al*., 2016), which could explain the absence of extracellular ROS production observed in response to *Pst*. Lipid composition and organization of the PM influence PM localization and stability of NLRs and consequently their activity in cell death inducing (Saile *et al*., 2021). By disturbing the PM organization, exogenous addition of d18:0 could also influence NLR localization and function.

Collectively, our results demonstrated that d18:0 could have a dual effect on the PM when co-infiltrated with *Pst AvrRpm1* by disturbing the organization and mechanical properties of the PM and the myristoylation process, and thus the redirection of effectors to the PM. Both actions prevent the interactions of NLR with its effector and subsequent defense responses.

## MATERIAL AND METHODS

### Plant material and growth conditions

Seeds of the *Arabidopsis* T-DNA insertion mutants *rpm1-3* (CS68739), *rpm1-rps2-rin4* (CS68760), and transgenic line *RPM1-myc rpm1* (CS68778) were obtained from the Nottingham *Arabidopsis* Stock Center (http://Arabidopsis.info). Mutants, transgenic lines and wild-type seeds Columbia-0 (Col-0) were grown in soil under 12 h-light/12 h-dark conditions (150 µmol/m^2^/s, 20°C, and 60% humidity) for 5 weeks.

### Chemicals

Sphinganine (d18:0), palmitoyl-2-linoleoyl-*sn*-glycero-3-phosphocholine (PLPC), sitosterol (sito), *D*-glucosyl-ß-1,1′-*N*-palmitoyl-*D*-erythro-sphingosine (d18:1/16:0 - GluCer or Glucosylceramide) were purchased from Avanti Polar Lipids. Sphingolipids were dissolved in ethanol (100%); PLPC, sitosterol and GluCer were dissolved in chloroform/methanol (2:1, v:v), and used without further purification. Protamine sulphate was purchased from Sigma-Aldrich (France).

### Bacterial cultures and inoculations

*Pst* DC3000 strains transformed to express the effectors AvrRpm1 (*Pst AvrRpm1*), AvrRpt2 (*Pst AvrRpt2*), AvrRps4 (*Pst AvrRps4*), AvrB (*Pst AvrB*) or AvrPphB (*Pst AvrPphB*) were provided by Prof. Jeff Dangl (University of North Carolina, USA) and Dr. Farid El-Kasmi (University of Tübingen, Germany), Prof. Brian Staskawicz (University of California, USA), Prof. Jane Parker (Max-Planck Institute, Cologne, Germany) and Dr. Brad Day (Michigan State university, USA), respectively. Bacteria were cultured overnight under agitation (180 rpm) at 28°C in liquid King’s B medium, supplemented with rifampicin (50 µg/mL) and kanamycin (50 µg/mL). Bacterial cells were collected by centrifugation, washed twice, and resuspended in 10 mM MgCl2. Bacterial solutions were infiltrated at 10^7^ CFU/mL on the abaxial side of *Arabidopsis* leaves using a 1 mL syringe without needle. Co-treatment of bacteria with 100 µM d18:0 was performed likewise. Control inoculations were conducted with 10 mM MgCl2. At least four leaves per plant were treated for each condition, with a minimum of three different plants.

### *In vitro* antibacterial assays

Bacteria were cultured as previously described before being distributed in 96 wells plates at 10^7^ CFU/mL in King’s B medium supplemented with appropriate antibiotics, under agitation and at 28°C. Bacteria were plated either alone, with ethanol (0.1%, v/v) or with d18:0 (100 µM). OD600 measurements were performed using the TECAN Spark® (USA) absorbance reader, every hour for 50 h. The presented data are obtained from growth curve experiments, from which the growth rate was extracted.

### Pathogen assay *in planta*

For bacterial count experiments, bacterial cells were cultured and inoculated as described. Four inoculated leaves were weighed and ground in 1 mL of 10 mM MgCl2 using a mortar and pestle. Appropriate dilutions were plated on King’s B medium with suitable antibiotics, and bacterial colonies were counted. Data are reported as means ±SD of the log (CFU/mg fresh weight) of three replicates.

### RNA extraction and qRT-PCR

Isolation of total RNA and real-time PCR were performed as described by Magnin-Robert et al. (2015). Gene-specific primers are available in the Supplemental Table S1. For each experiment, PCR was performed in triplicates. Transcript levels were normalized against those of the UBIQUITIN5 (*UBQ5*) gene, used as an internal control. Fold induction compared with untreated sample was calculated using the 2^-ΔΔCt^ method, where ΔΔCt= (CtGI[unknown sample]-CtGI[reference sample])-(CtUBQ5[unknown sample]-CtUBQ5[reference sample]) where GI is the gene of interest.

### Immunoblotting assay

Total proteins from pre-treated leaves of *RPM1-myc rpm1* plants collected 0, 2, 6 and 24 hpi were extracted and western-blotted according to Luzuriaga-Loaiza *et al*. (2018). Membrane hybridization was carried out through overnight incubation with recombinant monoclonal Anti-c-myc epitope tag [Clone 9E10] Mouse IgG1 kappa (Cliniscience, 1:1000) at 4°C, followed by a 1 h incubation with anti-mouse IgG HRP-conjugated secondary antibodies (Bio-Rad, 1:3000) at room temperature.

### Electrolyte leakage

Measurement of electrolyte leakage were performed as described in Magnin-Robert *et al*. (2015), except for leaf discs placed in each well of a 24-well plate containing 1 mL of distilled water. Conductivity measurements were then conducted over time using a B-771 LaquaTwin (Horiba) conductivity meter.

### Extracellular ROS production

Measurements of extracellular ROS production were performed as described in Magnin-Robert *et al*. (2015). Bacteria were added to a final concentration of 10^8^ or 10^9^ CFU/mL, as specified further. In tests involving d18:0, the molecule was added at a final concentration of 100 µM concomitantly with the elicitation solution.

### Phytohormone and camalexin quantifications

The quantification of SA, JA-isoleucine (JA-Ile) and camalexin was performed at 0 and 48 hpi by UPLC-nano-ESI-MS/MS as previously described in Herrfurth *et al*. (2020). Phytohormones were extracted with methyl-tert-butyl ether (MTBE), reversed phase-separated using an ACQUITY UPLC® system (Waters Corp., USA) and analysed by nanoelectrospray ionization (nanoESI) (TriVersa Nanomate^®^; Advion BioSciences, USA) coupled with an AB Sciex 4000 QTRAP^®^ tandem mass spectrometer (AB Sciex, USA) employed in scheduled multiple reaction monitoring mode. For quantification, 10 ng D4-SA (C/D/N Isotopes Inc., Canada), 20 ng D5-IAA (Eurisotop, Germany), and 10 ng D3-JA-Leu (kindly provided by Otto Miersch, Halle/Saale, Germany) were added at the beginning of the extraction procedure. For detection, the following mass transitions were used: 137/93 (declustering potential (DP) −25 V, entrance potential (EP) −6 V, collision energy (CE) −20 V) for SA, 141/97 (DP −25 V, EP −6 V, CE −22 V) for D4-SA, 179/135 (DP −35 V, EP −9 V, CE −14 V) for D5-indole-acetic acid, 199/141 (DP −60 V, EP −4 V, CE −32 V) for camalexin, 322/130 (DP −45 V, EP −5 V, CE −28 V) for JA-Ile and 325/133 (DP −65 V, EP −4 V, CE −30 V) for D3-JA-Leu.

### GUS reporter assays

To determine induction of *PR1*, GUS enzyme activity of *PR1::GUS Arabidopsis* plants was determined histochemically on leaves of the reporter line, treated as previously described, collected at 48 hpi, and placed in the GUS staining solution (50 mM sodium phosphate buffer - pH 7, 10mM EDTA, 1 mM K3Fe(CN)6, 2 mM X-GlcA, and 0,1% Triton X-100). The leaves were vacuum-infiltrated for 5 min and incubated for 4 h at 37°C. Next, they were washed with Milli-Q water and left overnight at 4°C in a discoloration / fixation solution (ethanol:acetic acid, 3:1). Once the discoloration process was complete, leaves were placed in 95% (v/v) ethanol.

### Lipid composition for plant biomimetic membranes

To mimic the lipid composition of plant PM, PLPC, β-sitosterol and GluCer were used, either in a mixture (60:20:20, molar ratio) or alone, depending on the experiment. This lipid composition was previously used to mimic plant PM (Deleu *et al*., 2014, 2019; Deboever *et al*., 2022).

### Adsorption of sphinganine onto lipid monolayers using Langmuir trough

Adsorption experiments at constant surface area were performed in a KSV (Helsinki, Finland) Minitrough (190 cm^3^) equipped with a Wilhelmy plate, as previously described (Nasir *et al*., 2016; Deleu *et al*., 2019). The maximal insertion pressure (MIP) and the differential П0 (dП0) were determined as previously described (Deboever *et al*., 2020).

### Preparation of large unilamellar vesicles

PLPC/ β-sitosterol/GluCer (60:20:20 molar ratio) large unilamellar vesicles (LUVs) were prepared as described previously (Lebecque et al., 2019). Tris-HCl buffer (10 mM, pH 7.4) was used for lipid film hydration and dispersion.

Extrusion was done *via* 13 passages through a 0.1 µM membrane filter using Liposofast extruder (AVESTIN®). Diameter and stability of the LUVs were confirmed by dynamic light scattering (DLS) measurements.

### Thermodynamic parameters of sphinganine interaction with lipid bilayers by isothermal titration calorimetry

For isothermal titration calorimetry (ITC) measurements, LUVs were prepared as described and resuspended to reach a final concentration of 1 mM in Tris-HCl buffer (10 mM, pH 7.4). Experiments were performed on a A VP-ITC microcalorimeter (Microcal Inc., Northampton, MA, USA) as previously described (Lebecque *et al*., 2018). Details can be found in supporting information section (Supplemental Material S1).

### Laurdan polarization on lipid vesicles LUVs

LUVs suspension at a concentration of 100 µM in lipids was incubated with 1 µM of Laurdan (Sigma-Aldrich) for 1h15. Then, wells of black 96-well microplates (Greiner Bio-One™ CellStar™, Fischer Scientific) were loaded with 100 µL of liposome solution per well. The fluorescence was recorded using Spark® microplate reader (Tecan) with an excitation wavelength at 360 (±35) nm and emission wavelengths at 430 (±20) nm and 485 (±20) nm. The fluorescence was recorded once before treatment and every 5 min for up to 20 minutes, after the addition of 100 µL of 2-times concentrated treatment.

The generalized polarization (GP) was defined as 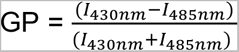, where I430nm and I485nm represents the blank-subtracted fluorescence intensities at emission wavelengths of 430 nm and 485 nm respectively. Variation of GP (ΔGP) is defined as the subtraction of GP measured at each time point following treatment from GP measured before treatment.

### Propensity of sphinganine to insert into a bilayer determined by the Impala procedure

The most energetically favorable position and orientation of d18:0 in an implicite lipid bilayer was determined with the same procedure described in Ducarme *et al*. (1998) and Franche *et al*. (2020). For more details, see Supplementary Material S1.

### Interaction energies between sphinganine and lipids determined by the Hypermatrix docking method

The hypermatrix docking method was employed, as previously described (Deleu *et al*., 2014; Deleu *et al*., 2019) to dock a molecule of sphinganine to a lipid system composed of either plant or bacterial lipids.

### Modeling of sphinganine/plant lipids monolayer interactions with Big Monolayer method

The big monolayer method, as described by Deleu *et al*. (2019), was used to visualize lipid domains. Three repetitions of the system were calculated. The simulations were performed using plant lipid mixture with or without d18:0 (88% lipids, 12% d18:0).

### Protoplast isolation

The isolation of mesophyll protoplasts was carried out by an enzymatic method as previously described (Yoo *et al*., 2007). The protoplasts were resuspended at 10^6^ protoplasts/mL in a storage solution (20 mM MES, 400 mM mannitol, 20 mM KCl, 10 mM CaCl2) until further use.

### Atomic force microscopy

Protoplasts were adhered to the bottom of a glass-bottom Petrie dish by coating with positive-charge-bearing protamine sulphate 1 % in water: the bottom of the Petri dish was covered with a thin layer of this solution during 2 min, rinsed 3 times with the protoplast storage solution and the remaining solution was discarded. Protoplast suspension was incubated for 5 min with final concentration 2 µM or 5 µM of d18:0 or control ethanol in protoplasts storage solution for 5 min. Then, pre-treated or not protoplasts suspension was gently deposited to the coating Petri dish, and the protoplasts were settled for 5 min.

QNM-LC-CAL probes (Bruker, Billerica, USA), having a resonance frequency of 45 kHz, a tip radius of 65 nm, a tip height of 16 µm and a nominal spring constant below 100 mN/m were calibrated on a non-compliant sample to extract the deflection sensitivity which turned out to be around 10 nm/V. Afterwards the tip was pulled off (of 100 Um) and tuned in liquid to determine the spring constant.

Optics (Nikon Ti 2000, Nikon, Tokyo, Japan) were used in bright field mode to browse through all protoplasts and focus on those having a 100 µm diameter. Therefore, the tip was engaged at the very center of each and forces curves were captured in single force spectroscopy mode. The atomic force microscopy (AFM) used was a Bioscope Catalyst (Bruker, Billerica, USA). A z ramp output was used, with a ramp size of 5 µm, a ramp rate of 0.25 Hz, a contact time of 50 msec, and a forward and backward velocities of 5 µm/sec. Each force curves contained 2048 points. The Poisson’s ratio was arbitrary set at 0.3. We used a relative trigger mode, a tip threshold of 30 nm and plotted the deflection error as data type. It was verified that there was no apparent change in diameter or x,y motion of the object during the acquisition.

The Young’s modulus was extracted according to the Hert’s theory and normalized as a function of the protoplasts’ volumes. A minimum of 50 curves per condition were recorded and experiments were repeated three times.

### Confocal microscopy

To determine the membrane order, protoplasts were labelled with 300 nM final concentration of fluorescent probe di-4-ANEPPDHQ (Invitrogen, stock solution 1.5 mM in DMSO) in protoplast storage solution, during 5 min. The labelled protoplasts suspension was gently deposited to the bottom of a glass bottom Petri dish and the protoplasts were settled for 5 min. The labelled protoplast suspension was incubated for 5 min with final concentration 2 µM or 5 µM of d18:0 or control ethanol for 5 min.

Fluorescence images were acquired using a laser scanning microscope LSM 880 from ZEISS (Carl ZEISS SAS, Rueil Malmaison, France) through 63x (ON 1.4) oil immersion objective with ZEN software. Di-4-ANEPPDHQ excitation was done at 488 nm and emission intensities were acquired between 540–560 nm (green image) and between 630–650 nm (red image). Ratiometric imaging was performed using the Fiji software (Schindelin *et al*., 2021).

## ACKNOWLEDGEMENT

The authors thank Prof. Jeff Dangl (University of North Carolina, USA), Dr. Farid El-Kasmi (University of Tübingen, Germany), Prof. Brian Staskawicz (University of California, USA), Prof. Jane Parker (Max-Planck Institute, Cologne, Germany) and Dr. Brad Day (Michigan State university, USA) for generously providing the different bacterial strains used in this study. This work was supported by the National Fund for Scientific Research (FNRS, Belgium), the SFR Condorcet (18ARC107), and the “Fondation Universitaire de Belgique” (Belgian University Foundation). Some of the experiments were carried out within the Nanomat platform (www.nanomat.eu) supported by the Ministère de l’Enseignement Supérieur et de la Recherche, the Région Grand Est, and FEDER funds from the European Community. GG was supported by the Foundation for Training in Industrial and Agricultural Research (FRIA, FNRS, grant 1.E.069.20F), and MD is Senior Research Associates of the FNRS. I.F. acknowledges funding by the Deutsche Forschungsgemeinschaft (INST 186/822-1).

## AUTHOR CONTRIBUTIONS

SDC and MD designed the research. EH, SV, AB, CT, CCh, CH and GG performed the research. EH, MD, SDC, FB analyzed and discussed the data. SC, SD, JC, CJ, FF, CCl, IF provided critical feedback on the manuscript. EH, MD and SDC wrote the paper and incorporated the input of the rest of the authors.

## SUPPLEMENTAL DATA

**Supplemental Figure S1.** Adsorption behaviour of d18:0 into lipid monolayer mimicking plant plasma membrane

**Supplemental Table S1.** Gene-specific primers used in real-time reverse transcription PCR.

**Supplementary Material S1**. Methods in biophysics.

