## Supplemental Figure S1 for "Dual action of sphinganine in the plant disease resistance to bacteria"

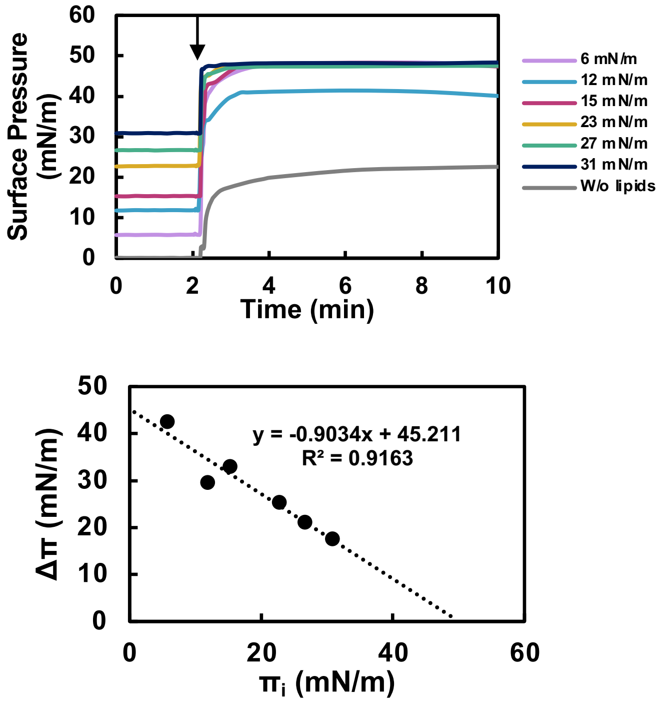


**FIGURE S1 Adsorption behaviour of d18:0 into lipid monolayer mimicking plant plasma membrane.**

(A) Kinetics of d18:0 (1.25 µM in the subphase Tris HCl, 10 mM, pH 7.4) adsorption to a bare interface or to the lipid monolayer (here the example of PLPC monolayer) at different initial surface pressures ($\Pi_i$). The arrow indicates the time at which d18:0 was injected underneath the bare interface or the PLPC monolayer. (B) Surface pressure variation following adsorption of d18:0 versus the initial surface pressure ($\Pi_i$) of the PLPC monolayer.
