## Supplementary Table S1 for "Dual action of sphinganine in the plant disease resistance to bacteria"

**Table S1:** Gene-specific primers used in real time reverse-transcription polymerase chain reaction.

| Gene | Forward Primer | Reverse primer | GenBank accession number |
|---|---|---|---|
| *AtPR1* | AACTACGCTGCGAACACGTG | TCACTTTGGCACATCCGAGTC | NM_127025 |
| *AtVSP1* | GGATCGAAGTTGACGCAAGTG | CTCAACCAAATCAGCCCATTG | NM_001125801 |
| *AtNMT1* | TCCTTCTGTTTACGAGTGGACGACATGT | CTCCAATATGCCAGCTCTGGTAATAACC | AF250956 |
| *AtUBQ5* | GGAAGAAGAAGACTTACACC | AGTCCACACTTACCACAGTA | NM_116090 |
