## Supplementary material S1 for "Dual action of sphinganine in the plant disease resistance to bacteria"

### SUPPLEMENTARY MATERIAL AND METHODS

#### Adsorption of sphinganine onto lipid monolayers using Langmuir trough

Lipid solutions were spread at the air-subphase interface to reach the desired initial surface pressure. After a 15 min wait for solvent evaporation and film stabilization, d18:0 was injected in the subphase (10 mM Tris buffer, pH 7.4), underneath the pre-formed lipid monolayer to a final concentration of 1.25 µM. The adsorption of d18:0 to the lipid monolayer was monitored by the increase of surface pressure as previously described (Nasir *et al*., 2016; Deleu *et al*., 2019). The injection of ethanol (0.1%) was used as control.

The MIP was obtained by linear regression of the plot $\Delta\Pi$ *vs* $\Pi_i$ and $d\Pi_0$ was calculated as follows: $d\Pi_0 = \Delta\Pi_0 - \Pi_e$

$\Delta\Pi_0$ corresponds to the y-intercept of the linear regression of the plot, and $\Pi_e$ is the surface pressure of d18:0 at the equilibrium when there is no lipid at the interface.

#### Thermodynamic parameters of sphinganine interaction with lipid bilayers by isothermal titration calorimetry

d18:0 was dissolved in DMSO (100%). All experiments were performed at 26°C. The reference cell was filled with degassed Milli-Q water. The sample cell (V= 1.4565 mL) was filled either with buffer (blank - Tris-HCl, 10 mM, pH 7.4), buffer + d18:0 (at a final concentration of 20 µM) or buffer + DMSO (control – 0.01%). This cell was continuously stirred at 305 rpm. The syringe (V= 300 µL) was filled with the LUV suspension + DMSO (1 mM in buffer + 0.01%). Both the LUV solution and the content of the sample cell were degassed by ultrasonication before use. The first injection of LUVs suspension was 2 µL and was not used in data analysis. Then, every 600 s, 10 µL of the suspension were injected in the sample cell. ITC data were analyzed using Origin 7.0 (Microcal) software following a previously described method (Heerklotz and Seelig, 2000; Lebecque *et al*., 2018).

**Propensity of sphinganine to insert into a bilayer determined by the Impala procedure**

First, the structure of d18:0 was constructed using HyperChem software (Hypercube, Inc.). The molecular geometry was optimized by systematic analysis of the torsion angles using the structure tree method as previously described (Lins *et al*., 1995). The most probable structure, corresponding to the lowest conformational energy, was used. The Impala procedure uses a Monte Carlo approach to simulate the insertion of d18:0 in an implicit lipid bilayer, as previously described (Ducarme *et al*., 1998; Franche *et al*., 2020). Briefly, this method employs an empirical forcefield to depict membrane physicochemical features while taking into account two types of restraints: the hydrophobic effect and lipid disturbances. The Z axis is assumed to be the only variable in membrane properties. The two restraints were calculated at each place by moving the d18:0 molecules along the Z axis by 1 Å steps from one side of the implicit membrane to the other and were later added to determine the total energy restraint.

**Interaction energies between sphinganine and lipids determined by the Hypermatrix docking method**

A molecule of d18:0 is fixed at the center of the system and oriented at the hydrophilic/hydrophobic interface. The lipid molecules, which are also oriented at the interface, are positioned around the center molecule and over $10^7$ positions (translations and rotations) are tested. Interactions such as Van der Waals, electrostatic, and hydrophobic are all taken into account when calculating the interaction energies. The energy of interaction along the coordinates of all assembly are gathered in a matrix and sorted according to decreasing values. The assembly with the lowest energy value is considered the energetically most stable.

**Modeling of sphinganine/plant lipids monolayer interactions with Big Monolayer method**

The initial stage consisted in aligning each pair of molecules in the system at the hydrophilic/hydrophobic interface according to Hypermatrix. Then, their interaction energies were computed, considering Van der Waals, electrostatic, and hydrophobic interactions. The second phase involved creating a grid of 200 × 200 molecules and utilizing a Monte Carlo procedure to minimize the system using the interaction energies computed in the previous step. Each molecule is represented by a pixel, producing an image of the molecular domains formed.
